# P53 TAD2 Domain (38-61) Forms Amyloid-like Aggregates in Isolation

**DOI:** 10.1101/2021.04.09.439126

**Authors:** Kundlik Gadhave, Shivani K Kapuganti, Pushpendra Mani Mishra, Rajanish Giri

**Author notes:** **Correspondence to**: Dr. Rajanish Giri, School of Basic Sciences, Indian Institute of Technology Mandi, Himachal Pradesh 175005, India.

## Abstract

In many cases, when cellular machinery is unable to restore changed protein conformations, they start sticking through exposed hydrophobic patches and form aggregates. A strong association between protein aggregation and Human diseases (such as Alzheimer’s, Parkinson’s, and Huntington’s disease) is well proven. p53 is a transcription factor that is also known as the guardian of the genome associated with cellular processes such as DNA repair, apoptosis, senescence, control of cell cycle, stress signaling and cellular homeostasis. The loss of function mutations in p53 have been implicated in several cancers. Experimental evidences have proposed a possible link between cancer and protein aggregation in evidence of the implication of amyloidogenic mutant proteins in ten different types of cancer. Aggregation studies focusing on different P53 domains, mostly, the central core domain and its mutants under the influence of various environmental conditions and P53 TAD domain (1-63) have been reported. P53 TADs interact with diverse cellular factors to modulate the function of P53 and elicit appropriate cellular response under different stress conditions. In this study, the aggregation of P53 TAD2 domain (38-61) have been studied in isolation. The aggregates were generated in-vitro in acidic pH conditions after in-silico scoring for amyloidogenic propensity and characterized using dye-based assays (ThT and bis-ANS fluorescence), CD spectroscopy, and microscopy (SEM, TEM and AFM). It was observed that P53 TAD2 follows nucleation-dependent kinetics and forms amyloid-like aggregates. On reductionists approach, this study highlights the nature of P53 TAD2 domain (amino acids 38-61) aggregation.

## Introduction

P53 is a transcription factor which under cellular stress conditions activates genes for cell growth arrest, DNA repair, apoptosis and senescence and suppresses genes involved in cell cycle progression and apoptosis inhibition [1]. The tertiary structure of P53 contains three main domains: an acidic N-terminal domain consisting of the transactivation domains (residues: 1-61) and a proline rich domain which maintains stability, a central core DNA binding domain (102-292) and a C-terminal domain consisting of the tetramerization domain (324-355) which is flanked by a nuclear localization signal and a nuclear export signal and a regulatory domain (363-393) [2,3]. The transactivation domain (residues: 1-61) is devided into two sub domains named TAD1 and TAD2.

Intrinsically disordered nature of the TAD domains has been established by CD and NMR analysis [4,5]. Even though P53 mutants have been associated with around fifty percent of human cancers, non-functional WT P53 also has been associated with a few cancers such as colon, breast cancers, neuroblastoma and retinoblastoma [6]. This could be due to loss of protein stability, misfolding and aggregation. Nuclear and cytoplasmic accumulation of P53 aggregates have been observed in such cancers [6]. Whereas a few studies have associated aggregation of P53 mutants with cancers such as breast, skin and prostate cancers [7–9]. Many human diseases such as Alzheimer’s, Parkinson’s, certain neuropathies, prion disease are caused due to protein misfolding and subsequent amyloid aggregation which disturbs the protein homeostasis by interfering with protein degradation system. Studies have demonstrated a dominant negative oncogenic activity of P53 resulting from similar amyloid aggregation of P53 mutants [10,11].

Misfolding of protein happen due to incorrect folding [12,13] and the most probable destiny for these misfolded proteins is self-aggregation [14]. After misfolding, protein may aggregate due to exposure of buried hydrophobic patches and its sticking with hydrophobic regions of other proteins if cellular machinery unable to restore it [14–16]. Aggregates of misfolded protein form amyloids by the rapid association of monomeric and oligomeric fibrils through the mechanism of nucleation-based polymerization that includes the lag, exponential and Saturation phase [17–20].

Among cancer-causing genes, one of the most frequently mutated and widely studied genes is p53 [21,22]. It is recognized as a transcription factor that activates transcription of the gene and also binds to the specific DNA response elements [22]. Investigations indicate that tumor-related mutations of p53 lead to gain of function in addition to previously reported loss of function which is most often phenomenon [23]. Mutant p53 which follow the prion-like aggregation leads to gain of function, dominant-negative, and loss of function effects [24]. Research have indicated that both wild-type and mutant p53 in tumor models shown the aggregation behavior like the classical amyloidogenic proteins (such as α-synuclein and Aβ) [7]. Aggregation of wild type p53 leads to its loss of anticancer activity and mutant p53 aggregation demonstrates enhanced amyloidogenic behavior and accelerate their aggregation [25]. So far, ten different types of cancer have been reported that found to be associated with amyloidogenic p53 mutations, proposing the link between Cancer and p53 aggregation [25]. Aggregation inhibition of both mutations induced as well as inherent p53 may stabilize its functional conformation and determine and approved the innovative approaches for Treatment as well as prevention of Cancer [25].

P53 aggregation-prone derivative peptides and its different domains such as C-terminal, central and N-terminal TAD domains have investigated in details for the aggregation and related biophysical property in recent years. A few studies have demonstrated the use of small molecules and peptides in preventing in-vitro aggregation of P53 mutants [26–28]. Understanding the nature of aggregation may help in developing better clinical interventions. Although, to develop anticancer drug it is crucial to understand the aggregation of p53 regions in all aspects, to proceed in this direction we are investigating the aggregation propensity of subdomain 2 of TAD i.e. AD2 in isolation. We further determine the biophysical properties of aggregated TAD2 in isolation were investigated and the aggregates were generated and characterized using spectroscopic and microscopic techniques.

## Material & Methods

### Reagents

Ammonium molybdate (NH4)6Mo7O24), Thioflavin T (ThT) and bis-ANS were procured from Sigma Aldrich, St. Louis, MO, USA. TEM Grids were obtained from TED PELLA INC. USA. The chemically synthesized p53 TAD2 domain peptide (_38_QAMDDLMLSPDDIEQWFTEDPGPD_61_) was obtained with >90% purity from Thermo Scientific, USA. All other chemicals, unless otherwise specified, were the highest purity available from local sources.

### P53 TAD2 domain amyloidogenic Propensity by In Silico Analysis

p53 TAD2 amino acid sequence was used in amyloidogenic β-aggregation propensity prediction algorithms. FoldAmyloid [29] and MetAmyl [30] servers were used to identify amyloidogenic regions. Intrinsic solubility profile of amino acids of p53TAD2 based on the PVIS score obtained by Camsol intrinsic server [31,32] at different pH and outputs were reproduced from http://www.mvsoftware.ch.cam.ac.uk/index.php/login. All the predictions were supported by literature.

### Preparation of p53 TAD2 peptide aggregates

0.14 mg of p53 TAD2 peptide dissolve in 350 ul of 10mM glycine-HCl buffer of pH 3 to get final concentration of 144 μM and this solution incubated at temperature of 9°C with shaking speed of 1200 rpm on thermomixer.

### Thioflavin T dye binding Assay

Thioflavin T dye intercalates into the amyloid fibrils and illustrates the sharp increase in the fluorescence intensity [33]. ThT assay was done as described previously [34,35]. The stock solution of ThT (200 μM) was prepared in distilled water. Then aggregated peptide was mixed with ThT with final concentrations of ThT and peptide was adjusted as 20 μM and 50 μM, respectively. The solution was added to a 96 wells plate and ThT fluorescence intensity was measured on an infinite M200 PRO plate reader with excitation wavelength at 450 nm and emission wavelength at 490 nm, respectively.

### Bis-ANS binding assay

The aggregated p53 TAD2 peptide were evaluated for its ability to bind with bis-ANS dye, a dye which is commonly used to detect the exposed hydrophobic surfaces of proteins/peptides and their aggregates [36]. The p53 TAD2 aggregated sample (50 μM) were collected and mixed with the bis-ANS (30 μM). After incubated for 10 minutes at RT, samples were measured for fluorescence intensity by excitation at 380 nm, and emission was recorded between 414 and 650 nm. bis-ANS with buffer used as negative control.

### Circular Dichroism (CD) Spectroscopy

The secondary structure information of p53 TAD2 peptide amyloids was obtained from far-UV CD spectra recorded from 190 nm to 260 nm using JASCO J-1500 CD spectrophotometer. The aggregated samples were diluted in 10 mM Glycin-HCl buffer (pH-3) at a concentration of 20μM. The freshly dissolved p53 TAD2 peptide was used to record the spectra in monomeric form as a control experiment. The CD spectra were collected in 1mm path length cuvette at 25 °C with the scan speed of 50□nm/min, data pitch of 0.5 nm, a response time of 1s, and a bandwidth of 2□nm. Spectra for respective blank were subtracted from every sample.

### Field Emission Scanning Electron Microscopy (FE-SEM)

The FE-SEM imaging for p53 TAD2 peptide aggregates was performed using a Nova Nano SEM-450; FEI instrument. The 10 μl aggregated sample of p53 TAD2 peptide was mounted on pre-cleaned silicon wafers and then dried overnight. Then the sample was sputter coated with gold at a thickness of approximately 5 nm and observed under FE-SEM.

### High Resolution Transmission Electron Microscopy (HR-TEM)

For the morphological study, 10 μl of p53 TAD2 peptide aggregated sample was mounted on formvar-coated 200 mesh copper grids and incubated for 60 sec at RT. Further, grid were washed by gently spinning in a droplet of water for 30 sec. This process was repeated three times. The grids were negatively stained with 3% w/v aqueous solution of ammonium molybdate. The grids were then air-dried overnight, and images were recorded in HR-TEM (FP 5022/22-Tecnai G2 20 S-TWIN, FEI) with accelerating at 200 kV.

### Atomic Force Microscopy (AFM)

Aggregated p53 TAD2 peptide sample was 20-fold diluted with water and the 10 μL diluted sample was mounted on freshly cleaved mica sheet. After two minutes, the sample washed with deionized water and then allowed to dry in a desiccator for overnight. Finally, the imaging was performed on tapping-mode AFM (Dimension Icon from Bruker).

## Results & Discussion

### Results

#### In silico screening and prediction of appropriate pH condition

In silico screening of TAD2 peptide sequence with FoldAmyloid [29] and MetAmyl [30] server reveal the presence of aggregation-prone residues and regions. **Figure 1A** shows the aggregation prone region (residues 51-55) of TDA2 by FoldAmyloid and (**Figure 1B**) represents amloidogenic region (residues 50-55) by MetAmyl. CamSol intrinsic solubility [31,32] profile for each amino acid and combine protein variant intrinsic solubility score determined by CamSol Intrinsic server. Intrinsic solubility profiling of each amino acid shown in **Figure 1C**. The combined solubility score versus pH graph shown in **Figure 1D**, most suitable pH for aggregation experiment mark with asterisk.

**Figure 1.**
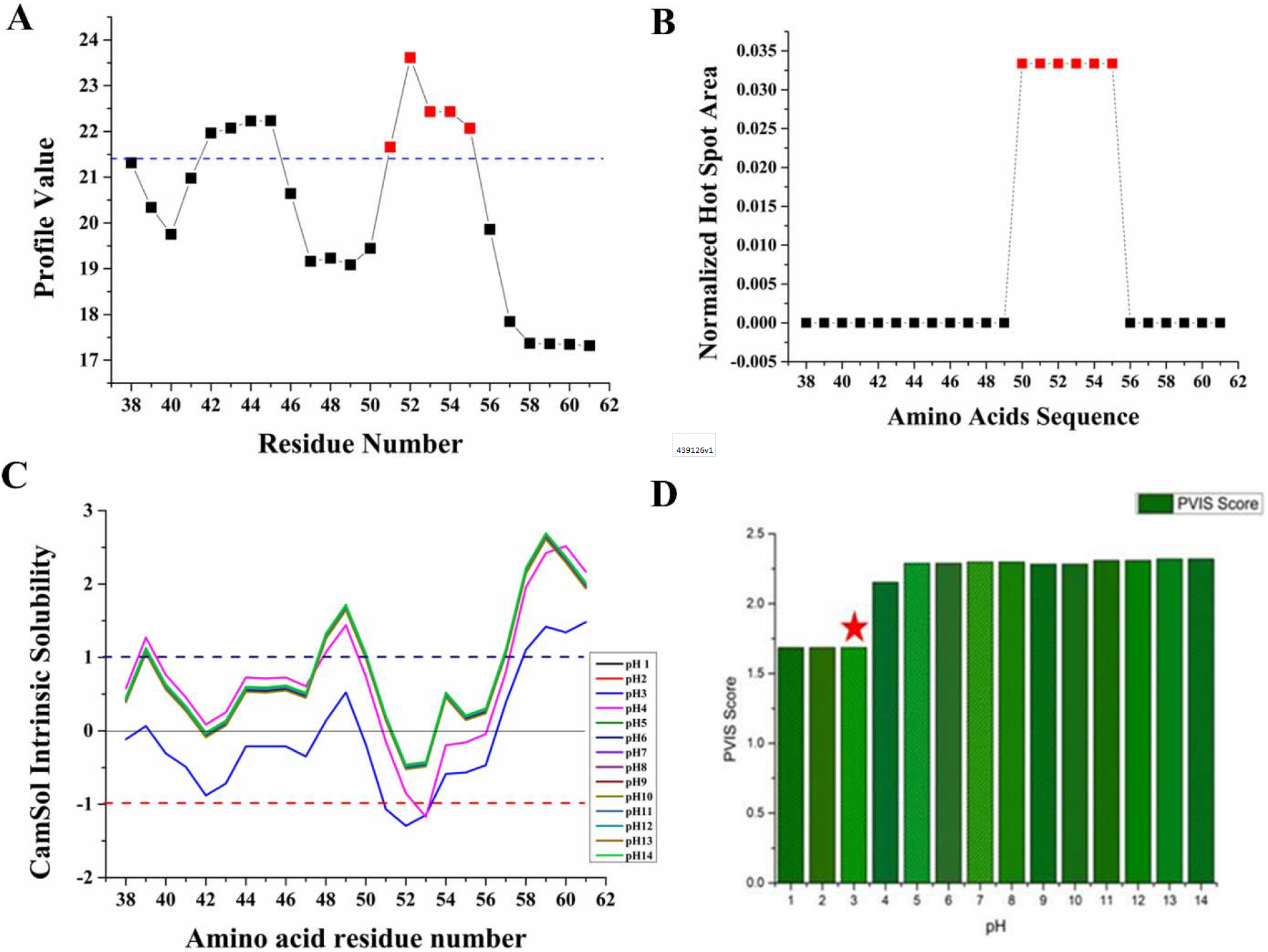
Prediction of p53 TAD2 aggregation propensity. (a) Aggregation prone regions using FoldAmyloid server. Residues above blue line are amyloidogenic. Region marked with red is amyloidogenic. (b) Aggregation prone regions using MetAmyl. Region marked with red is amyloidogenic. (c) Camsol Intrinsic solubility profile at different pH. (d) Prediction of best pH (red asterisked) for aggregation based on the PVIS score obtained by Camsol intrinsic server and supported by the literature.

#### p53 TAD2 aggregates are ThT and ANS-Positive

Thioflavin T dye used frequently for the characterization of formed amyloid from protein or peptide without showing significant interaction to corresponding monomeric peptide or protein [37]. Amyloid bonded dye when excited at wavelength 450 nm with scanning range 470 nm-700 nm; shows enhanced fluorescence at approximately 490 nm. **Figure 2A** shows the Thioflavin T binding assay for p53 TAD2, where red color fluorescence spectra positively indicate amyloid formation and black color spectra indicate negative control (ThT with buffer). **Figure 2B** represent ThT fluorescence kinetics for p53 TAD2 aggregation. Red color indicates p53 TAD2 and black color indicates ThT control samples at different time point. It shows fast aggregation of TAD2 with typical nucleation-dependent kinetic.

**Figure 2:**
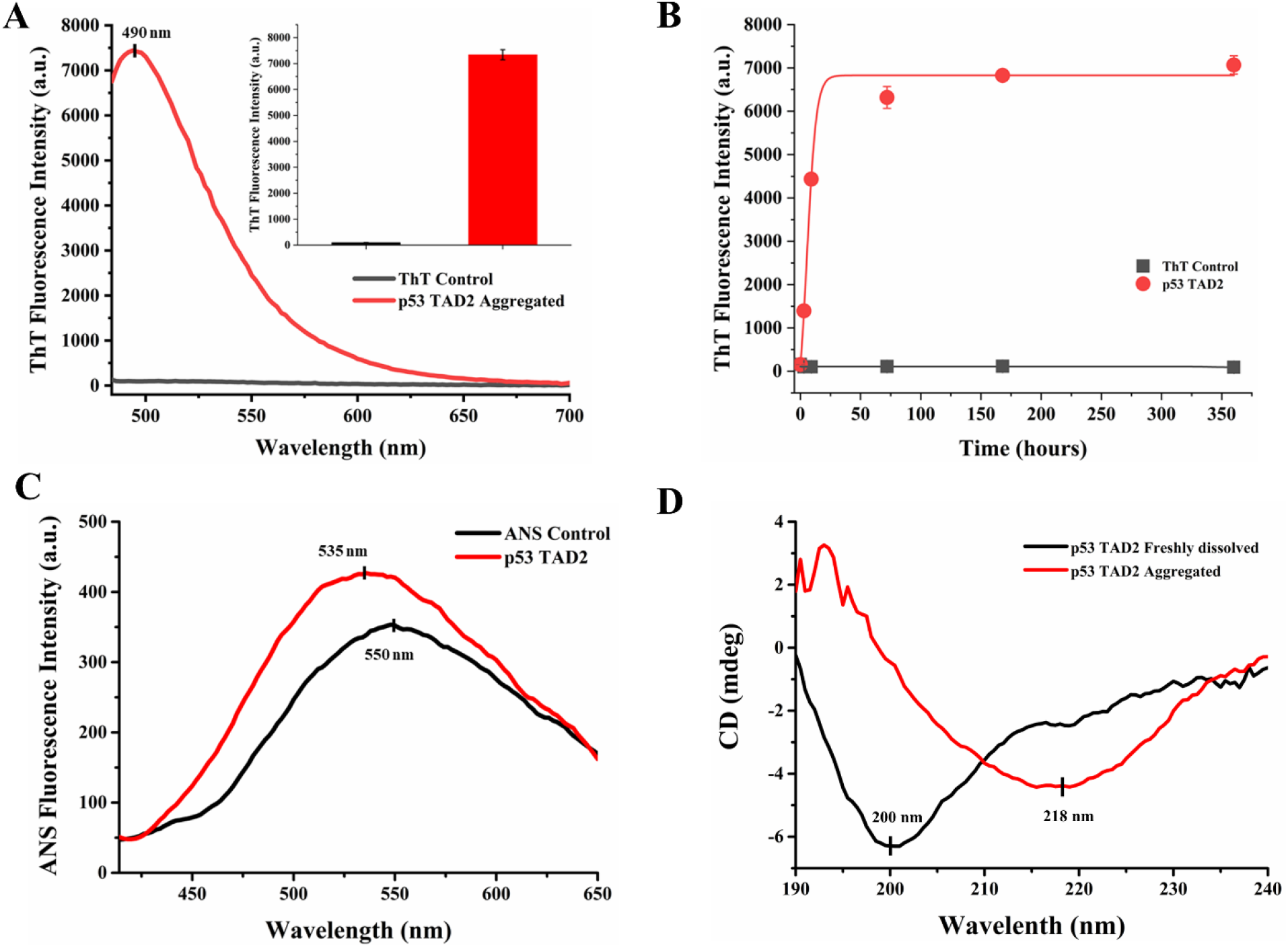
Dye binding assay and CD spectroscopy for p53 TAD2. (a) Thioflavin T fluorescence assay for p53TAD2, fluorescence intensity at 482 nm (red color) indicates the presence of amyloid aggregates, while black color spectra represent ThT dye. (b) ThT fluorescence kinetics. (c) bis-ANS assay for p53 TAD2, increase fluorescence with blue shift (red spectra) indicate the presence of amyloids in each sample, while black color spectra indicate fluorescence intensity of bis-ANS at pH 3. (d) CD Spectra for monomeric and aggregated p53 TAD2.

Bis-ANS is commonly used dye for the detection of exposed hydrophobic regions. It binds to an amyloid form of protein or peptide and enables it to characterize protein aggregation. A characteristic blue shift with increase in fluorescence intensity observed when bis-ANS binds to amyloid (**Figure 2 C**). The red color spectra show the blueshift at fluorescence maxima near to 535 nm, when ThT bound to TAD2 amyloid and excited at wavelength 380 nm with scan range between 414-650nm. Black color spectra indicate negative control (bis-ANS with buffer).

#### p53 TAD2 aggregates are rich in β-sheet structure

CD spectrum used to determine the configuration of p53 TAD2 peptide aggregates. The CD spectra of freshly dissolved p53 TAD2 peptide is considered as monomeric protein were used as negative control. The monomeric peptide at pH3 gives negative band near to 200 nm (**Figure 2D**), the characteristic spectra of disordered protein. This spectrum is already reported in previous study [38]. Furthermore, the CD spectra were recorded for aggregated p53 TAD (**Figure 2D**) shows the negative band near to 218 nm (characteristic point for well-defined β-conformation) that reveal the aggregates contain the major configuration of β-sheets.

#### Amyloid Fibrillar structure of p53 TAD2 visualized by TEM, SEM, and AFM

Matured fibrillar structure of formed p53 TAD2 amyloid shown by SEM, TEM, and AFM in **Figure 3; a, b, and c** respectively. SEM image demonstrates the rod-like robust appearance of formed aggregate, detail investigation by TEM reveals the fibrillar structured aggregates, and AFM image shows a tightly packed clump of the rod-like structure. Each technique investigates the details of the amyloid structure at different magnification and visualization level. The diameter of fibrils were determined from SEM images represents 20-30 nm.

**Figure 3:**
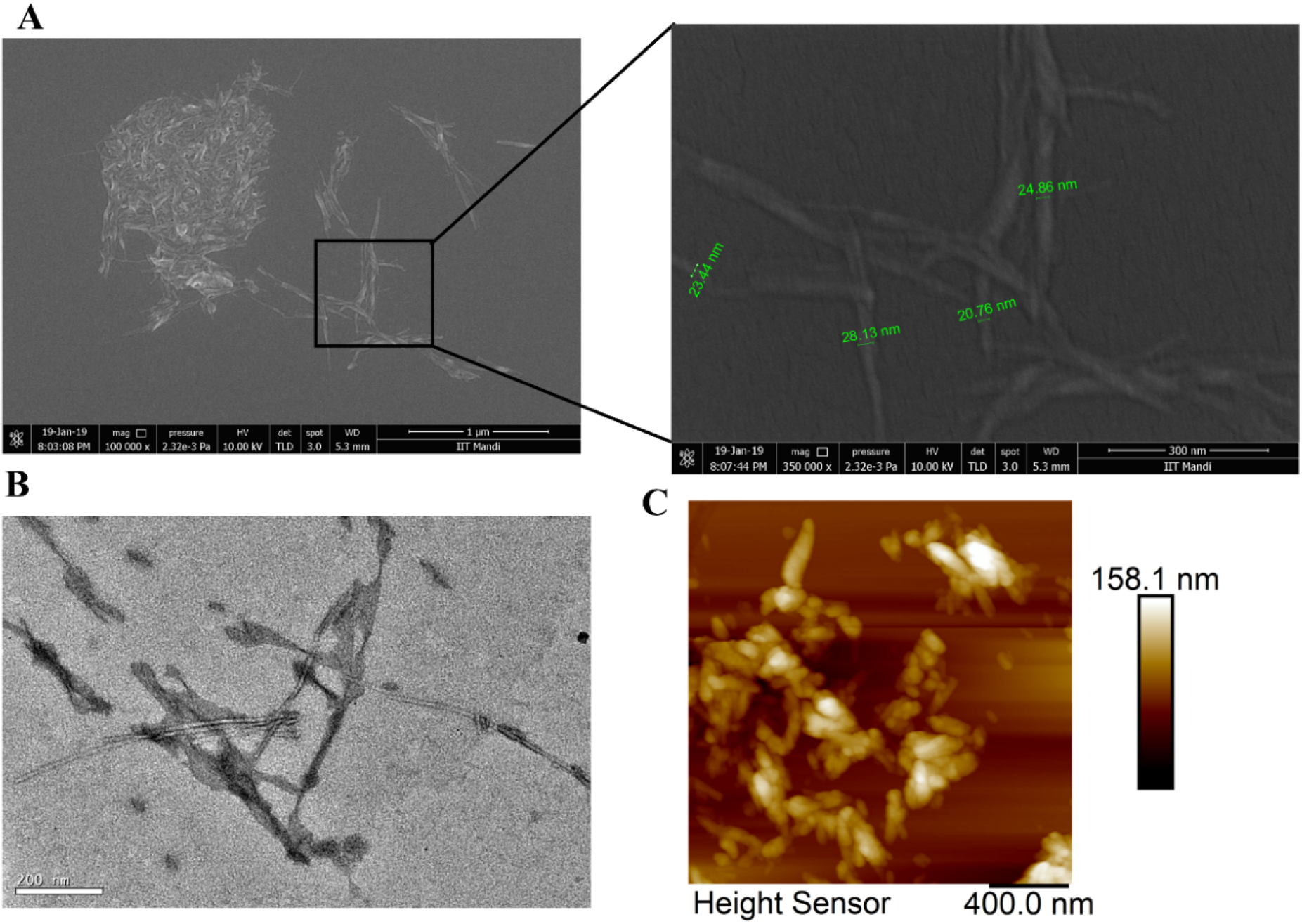
Morphological characterization of p53 TAD2 aggregates by microscopy. P53 TAD2 peptide were incubated at glycine buffer, pH3 for 15 days. Samples were prepared and visualized by (a) SEM. (b) TEM. and (c) AFM image for aggregated p53 TAD2.

## Discussion

The pathogenic protein aggregation is associated with various diseases such as AD, PD, and T2DM. The tumor suppressor p53 is also amyloidogenic and forms aggregates with morphology like classical amyloidogenic proteins, such as Aβ and α- synuclein. These results into loss of anticancer activity of p53 when it aggregates, suggesting a connection between p53 aggregation and cancer [25]. Interestingly, p53 aggregation has been observed in in vivo studies, patient tissues, and biopsies that show correlation between p53 aggregation with cancer. Aggregation of p53 leads to loss of its tumor-suppressor function, negative dominance effects and cancers with a worsened prognosis with tumor growth [11,39,40]. The aggregation of the entire p53 protein and its three functional domains into amyloid fibrils has been reported. Mutations in the p53 DNA-binding domain (p53C) form amyloid fibrils. Moreover, aggregation of mutant p53 is reported in breast cancer and malignant skin tumors [40].

Accumulated aggregates of WT and mutant p53 leading to loss of tumor suppression activity of P53 have been associated with some cancers. The TAD domains of P53 are responsible for interaction with various cellular proteins that modulate the transcriptional activity of P53. It has been demonstrated that the loss of phosphorylation sites (TAD1: S6, S9, S15, T18, S20, S33, S37, TAD2: S46, T55) and the hydrophobic sites (TAD1: L22, W23, TAD2: W53, F54) in these domains led to loss of binding capacity with other proteins [41]. TAD domains are intrinsically disordered that assume a secondary structure in binding with target proteins. Research has shown that this region assumes an alpha helical structure on binding to MDM2 and CBP [42,43]. Protein aggregation arises due to a disruption of this folding-unfolding transition equilibrium. Aggregation favors the formation of secondary structures that are not native at the same time preventing formation of tertiary structures. This can also result in the co-aggregation of P53 with other family proteins which also has been reported [10].

The proper folding of proteins is facilitated by chaperones. Mildly destabilizing conditions are known to lead to aggregation of acidic domains of proteins. Some compartments of the cells have mildly low pH. It has already been hypothesized that low pH near membranes can aid in beta sheet formations [44]. The aggregation of P53 associated with cancers may very well start with the TAD regions. Some mutations of TAD domains have been implicated in cancers as mentioned earlier. How these established mutations affect the conformation, folding and aggregation of these domains and the entire protein needs to be further investigated. Amyloidogenic mutations in p53 have reported in different types of cancer that reveal the strong relationship among the incidents of mutation, protein misfolding, protein aggregation and causation of cancer [25,40]. In addition to the mutations, various other factors responsible for the aggregation of protein and its induced toxicity and ultimately cell death and disease [25,40]. The motivation of current experimentation is to explore and analyze the effect and property of amyloid that could be formed due to various reasons as discussed above.

For this purpose, we have artificially induced the formation of amyloid in p53 TDA2 in Glycine-HCl buffer of pH-3 at Temperature 9°C for an incubation period of 15 days with constant shaking at 1200 rpm after scoring for amyloid aggregation propensity using CamSol server. Validation of formed amyloid has done with ThT and bis-ANS assay and CD spectroscopy, while morphological characterization has performed with microscopic techniques SEM, TEM, and AFM. ThT is a benzothiazole salt that binds to the β-sheet rich structure. The formation of TAD2 aggregates was confirmed using ThT fluorescence and ThT kinetics assay which gives a maximum emission peak at ~490 nm upon binding to β-sheets in fibrillar aggregates. bis-ANS also represents typical blue shift with increase in fluorescence. The β-sheet nature of formed aggregates was confirmed using CD spectroscopy. The fibrillar morphology characteristic to amyloid aggregation was confirmed using SEM, TEM and AFM. Identification of aggregation-prone regions within p53 and their mechanistic characterization would enable us to determine the therapeutic strategies and lead molecules that stabilize the functional form of protein and help in dissolving the already formed aggregates.

## Conclusion

Various factors including altered cellular mechanism and changed intracellular environment responsible for the aggregation of p53 protein. Upon aggregation, p53 loses its normal function and gain the cytotoxic function that leads to the disease. Our results revealed that the incubation of p53 TAD2 in acidic condition resulted in the folding of p53 TAD2 in β-sheet secondary structure. ThT fluorescence kinetics revealed that the p53 TAD2 follows nucleation-dependent aggregation with fast aggregation. CD spectroscopy results demonstrates the amyloid formation attended by the conformational change in p53 TAD2 from random coil to β-sheet rich structure. Microscopic images revealed the formation of typical amyloid-like aggregates with diameter ranging from 20-30 nm. This study provides useful information on the amyloidogenic nature of p53 TAD2 domain that will assist in the future to understand mechanism of full-length TAD aggregation and its involvement in the cancer pathogenesis.

## Author Contributions

RG: Conception, design, review, and writing of the manuscript. KG and SKK, and PMM performed experiments. KG, SKK, and PMM analyzed data and wrote the manuscript.

## Acknowledgments

Authors are grateful to Indian Institute of Technology Mandi (BioX and AMRC center) for all the facilities. This work was supported by the funding from Department of Biotechnology (DBT), Government of India (Project: BT/PR15453/BRB/10/1460/2015) to RG.

## References

[1] B.I. Batinac T, Gruber F, Lipozencić J, Zamolo-Koncar G, Stasić A, Protein p53-- structure, function, and possible therapeutic implications, Acta Dermatovenerol Croat. 11 (2003) 225–30. https://pubmed.ncbi.nlm.nih.gov/14670223/ (accessed March 16, 2021).

[2] M. Li, C.L. Brooks, F. Wu-Baer, D. Chen, R. Baer, W. Gu, Mono-Versus Polyubiquitination: Differential Control of p53 Fate by Mdm2, Science (80-.). 302 (2003) 1972–1975. doi:10.1126/science.1091362.

[3] S. Rigacci, M. Bucciantini, A. Relini, A. Pesce, A. Gliozzi, A. Berti, M. Stefani, The (1-63) region of the p53 transactivation domain aggregates in vitro into cytotoxic amyloid assemblies, Biophys. J. 94 (2008) 3635–3646. doi:10.1529/biophysj.107.122283.

[4] S. Bell, C. Klein, L. Müller, S. Hansen, J. Buchner, p53 contains large unstructured regions in its native state, J. Mol. Biol. 322 (2002) 917–927. doi:10.1016/S0022-2836(02)00848-3.

[5] M. Wells, H. Tidow, T.J. Rutherford, P. Markwick, M.R. Jensen, E. Mylonas, D.I. Svergun, M. Blackledge, A.R. Fersht, Structure of tumor suppressor p53 and its intrinsically disordered N-terminal transactivation domain, Proc. Natl. Acad. Sci. U. S. A. 105 (2008) 5762–5767. doi:10.1073/pnas.0801353105.

[6] U.M. Moll, A.G. Ostermeyer, R. Haladay, B. Winkfield, M. Frazier, G. Zambetti, Cytoplasmic sequestration of wild-type p53 protein impairs the G1 checkpoint after DNA damage., Mol. Cell. Biol. 16 (1996) 1126–1137. doi:10.1128/mcb.16.3.1126.

[7] A.P.D. Ano Bom, L.P. Rangel, D.C.F. Costa, G.A.P. De Oliveira, D. Sanches, C.A. Braga, L.M. Gava, C.H.I. Ramos, A.O.T. Cepeda, A.C. Stumbo, C. V. De Moura Gallo, Y. Cordeiros, J.L. Silva, Mutant p53 aggregates into prion-like amyloid oligomers and fibrils: Implications for cancer, J. Biol. Chem. 287 (2012) 28152–28162. doi:10.1074/jbc.M112.340638.

[8] C.B. Levy, A.C. Stumbo, A.P.D. Ano Bom, E.A. Portari, Y. Carneiro, J.L. Silva, C. V. De Moura-Gallo, Co-localization of mutant p53 and amyloid-like protein aggregates in breast tumors, Int. J. Biochem. Cell Biol. 43 (2011) 60–64. doi:10.1016/j.biocel.2010.10.017.

[9] M. Kluth, S. Harasimowicz, L. Burkhardt, K. Grupp, A. Krohn, K. Prien, J. Gjoni, T. Haß, R. Galal, M. Graefen, A. Haese, R. Simon, J. Hühne-Simon, C. Koop, J. Korbel, J. Weischenfeld, H. Huland, G. Sauter, A. Quaas, W. Wilczak, M.C. Tsourlakis, S. Minner, T. Schlomm, Clinical significance of different types of p53 gene alteration in surgically treated prostate cancer, Int. J. Cancer. 135 (2014) 1369–1380. doi:10.1002/ijc.28784.

[10] J. Xu, J. Reumers, J.R. Couceiro, F. De Smet, R. Gallardo, S. Rudyak, A. Cornelis, J. Rozenski, A. Zwolinska, J.C. Marine, D. Lambrechts, Y.A. Suh, F. Rousseau, J. Schymkowitz, Gain of function of mutant p53 by coaggregation with multiple tumor suppressors, Nat. Chem. Biol. 7 (2011) 285–295. doi:10.1038/nchembio.546.

[11] L.P. Rangel, D.C.F. Costa, T.C.R.G. Vieira, J.L. Silva, The aggregation of mutant p53 produces prion-like properties in cancer, Prion. 8 (2014). doi:10.4161/pri.27776.

[12] C. Soto, Protein misfolding and disease; protein refolding and therapy, FEBS Lett. 498 (2001) 204–207. doi:10.1016/S0014-5793(01)02486-3.

[13] J.W. Kelly, Alternative conformations of amyloidogenic proteins govern their behavior, Curr. Opin. Struct. Biol. 6 (1996) 11–17. doi:10.1016/S0959-440X(96)80089-3.

[14] I. Moreno-Gonzalez, C. Soto, Misfolded protein aggregates: Mechanisms, structures and potential for disease transmission, Semin. Cell Dev. Biol. 22 (2011) 482–487. doi:10.1016/j.semcdb.2011.04.002.

[15] M.J. Gething, J. Sambrook, Protein folding in the cell, Nature. 355 (1992) 33–45. doi:10.1038/355033a0.

[16] C.J. Roberts, Non-native protein aggregation kinetics, Biotechnol. Bioeng. 98 (2007) 927–938. doi:10.1002/bit.21627.

[17] J.W. Kelly, The alternative conformations of amyloidogenic proteins and their multi-step assembly pathways, Curr. Opin. Struct. Biol. 8 (1998) 101–106. doi:10.1016/S0959-440X(98)80016-X.

[18] H.A. Lashuel, B.M. Petre, J. Wall, M. Simon, R.J. Nowak, T. Walz, P.T. Lansbury, α-synuclein, especially the parkinson’s disease-associated mutants, forms pore-like annular and tubular protofibrils, J. Mol. Biol. 322 (2002) 1089–1102. doi:10.1016/S0022-2836(02)00735-0.

[19] M.A. Poirier, H. Li, J. Macosko, S. Cai, M. Amzel, C.A. Ross, Huntingtin spheroids and protofibrils as precursors in polyglutamine fibrilization, J. Biol. Chem. 277 (2002) 41032–41037. doi:10.1074/jbc.M205809200.

[20] S.I.A. Cohen, M. Vendruscolo, C.M. Dobson, T.P.J. Knowles, From macroscopic measurements to microscopic mechanisms of protein aggregation, J. Mol. Biol. 421 (2012) 160–171. doi:10.1016/j.jmb.2012.02.031.

[21] W.A. Freed-Pastor, C. Prives, Mutant p53: One name, many proteins, Genes Dev. 26 (2012) 1268–1286. doi:10.1101/gad.190678.112.

[22] J.T. Zilfou, S.W. Lowe, Tumor suppressive functions of p53., Cold Spring Harb. Perspect. Biol. 1 (2009). doi:10.1101/cshperspect.a001883.

[23] M. Oren, V. Rotter, Mutant p53 gain-of-function in cancer., Cold Spring Harb. Perspect. Biol. 2 (2010). doi:10.1101/cshperspect.a001107.

[24] G.A.P. de Oliveira, L.P. Rangel, D.C. Costa, J.L. Silva, Misfolding, aggregation, and disordered segments in c-Abl and p53 in human cancer, Front. Oncol. 5 (2015). doi:10.3389/fonc.2015.00097.

[25] H. Gong, X. Yang, Y. Zhao, R.B. Petersen, X. Liu, Y. Liu, K. Huang, Amyloidogenicity of p53: A Hidden Link Between Protein Misfolding and Cancer., Curr. Protein Pept. Sci. (2014). http://www.ncbi.nlm.nih.gov/pubmed/25440023 (accessed March 16, 2021).

[26] Z. Chen, J. Chen, V.G. Keshamouni, M. Kanapathipillai, Polyarginine and its analogues inhibit p53 mutant aggregation and cancer cell proliferation in vitro, Biochem. Biophys. Res. Commun. 489 (2017) 130–134. doi:10.1016/j.bbrc.2017.05.111.

[27] M. Kanapathipillai, Z. Chen, Inhibition of p53 Mutant Peptide Aggregation In Vitro by Cationic Osmolyte Acetylcholine Chloride, Protein Pept. Lett. 24 (2017) 353–357. doi:10.2174/0929866524666170123142858.

[28] A. Soragni, D.M. Janzen, L.M. Johnson, A.G. Lindgren, A. Thai-Quynh Nguyen, E. Tiourin, A.B. Soriaga, J. Lu, L. Jiang, K.F. Faull, M. Pellegrini, S. Memarzadeh, D.S. Eisenberg, A Designed Inhibitor of p53 Aggregation Rescues p53 Tumor Suppression in Ovarian Carcinomas, Cancer Cell. 29 (2016) 90–103. doi:10.1016/j.ccell.2015.12.002.

[29] S.O. Garbuzynskiy, M.Y. Lobanov, O. V. Galzitskaya, FoldAmyloid: A method of prediction of amyloidogenic regions from protein sequence, Bioinformatics. 26 (2009) 326–332. doi:10.1093/bioinformatics/btp691.

[30] M. Emily, A. Talvas, C. Delamarche, MetAmyl: A METa-predictor for AMYLoid proteins, PLoS One. 8 (2013). doi:10.1371/journal.pone.0079722.

[31] P. Sormanni, F.A. Aprile, M. Vendruscolo, The CamSol method of rational design of protein mutants with enhanced solubility, J. Mol. Biol. 427 (2015) 478–490. doi:10.1016/j.jmb.2014.09.026.

[32] P. Sormanni, L. Amery, S. Ekizoglou, M. Vendruscolo, B. Popovic, Rapid and accurate in silico solubility screening of a monoclonal antibody library, Sci. Rep. 7 (2017). doi:10.1038/s41598-017-07800-w.

[33] C. Xue, T.Y. Lin, D. Chang, Z. Guo, Thioflavin T as an amyloid dye: Fibril quantification, optimal concentration and effect on aggregation, R. Soc. Open Sci. 4 (2017). doi:10.1098/rsos.160696.

[34] K. Udit Saumya, K. Gadhave, A. Kumar, R. Giri, Zika Virus Capsid Anchor Forms Cytotoxic Amyloid-like Fibrils Schematic representation of Zika virus Capsid anchor forming amyloid aggregates with cytotoxic and hemolytic properties, BioRxiv. (2020). doi:10.1101/2020.11.13.381988.

[35] K. Gadhave, R. Giri, Amyloid formation by intrinsically disordered trans-activation domain of cMyb, Biochem. Biophys. Res. Commun. 524 (2020) 446–452. doi:10.1016/j.bbrc.2020.01.110.

[36] A. Hawe, M. Sutter, W. Jiskoot, Extrinsic fluorescent dyes as tools for protein characterization, Pharm. Res. 25 (2008) 1487–1499. doi:10.1007/s11095-007-9516-9.

[37] H. LeVine, Quantification of β-sheet amyloid fibril structures with thioflavin T, Methods Enzymol. 309 (1999) 274–284. doi:10.1016/S0076-6879(99)09020-5.

[38] D. Kumar, P.M. Mishra, K. Gadhave, R. Giri, Conformational dynamics of p53 N-terminal TAD2 region under different solvent conditions, Arch. Biochem. Biophys. 689 (2020). doi:10.1016/j.abb.2020.108459.

[39] M. Kanapathipillai, Treating p53 mutant aggregation-associated cancer, Cancers (Basel). 10 (2018). doi:10.3390/cancers10060154.

[40] D.C.F. Costa, G.A.P. de Oliveira, E.A. Cino, I.N. Soares, L.P. Rangel, J.L. Silva, Aggregation and prion-like properties of misfolded tumor suppressors: Is cancer a prion disease?, Cold Spring Harb. Perspect. Biol. 8 (2016). doi:10.1101/cshperspect.a023614.

[41] N. Raj, L.D. Attardi, The transactivation domains of the p53 protein, Cold Spring Harb. Perspect. Med. 7 (2017). doi:10.1101/cshperspect.a026047.

[42] B. Shan, D.W. Li, L. Brüschweiler-Li, R. Brüschweiler, Competitive binding between dynamic p53 transactivation subdomains to human MDM2 protein: Implications for regulating the p53·MDM2/MDMX interaction, J. Biol. Chem. 287 (2012) 30376–30384. doi:10.1074/jbc.M112.369793.

[43] C.W. Lee, M. Arai, M.A. Martinez-Yamout, H.J. Dyson, P.E. Wright, Mapping the interactions of the p53 transactivation domain with the KIX domain of CBP, Biochemistry. 48 (2009) 2115–2124. doi:10.1021/bi802055v.

[44] R. Li, Z. Wu, Y. Wangb, L. Ding, Y. Wang, Role of pH-induced structural change in protein aggregation in foam fractionation of bovine serum albumin, Biotechnol. Reports. 9 (2016) 46–52. doi:10.1016/j.btre.2016.01.002.

